# Rules of Contact Inhibition of Locomotion for Cells on Suspended Nanofibers

**DOI:** 10.1101/2020.05.28.122218

**Authors:** Jugroop Singh, Brian A. Camley, Amrinder S. Nain

## Abstract

Contact inhibition of locomotion (CIL), in which cells repolarize and move away from contact, is now established as a fundamental driving force in development, repair, and disease biology. Much of what we know of CIL stems from studies on 2D substrates that fail to provide an essential biophysical cue – the curvature of extracellular matrix fibers. We discover rules controlling outcomes of cell-cell collisions on suspended nanofibers, and show them to be profoundly different from the stereotyped CIL behavior known on 2D substrates. Two approaching cells attached to a single fiber do not repolarize upon contact but rather usually migrate past one another. Fiber geometry modulates this behavior: when cells are attached to two fibers, reducing their freedom to reorient, only one of a pair of colliding cells repolarizes on contact, leading to the cell pair migrating as a single unit. CIL outcomes also change when one cell has recently divided and moves with high speed– cells more frequently walk past each other. In collisions with division in the two-fiber geometry, we also capture rare events where a daughter cell pushes the non-dividing cell along the fibers. Our computational model of CIL in fiber geometries reproduces the core qualitative results of the experiments robustly to model parameters. Our model shows that the increased speed of post-division cells may be sufficient to explain their increased walk-past rate. Our results suggest that characterizing cell-cell interactions on flat substrates, channels, or micropatterns is not sufficient to predict interactions in a matrix – the geometry of the fiber can generate entirely new behaviors.

**Significance:** When cells heal a wound or invade a new area, they coordinate their motion. Coordination is often studied by looking at what happens after pairs of cells collide. Post-collision, cells often exhibit contact inhibition of locomotion– they turn around and crawl away from the point where they touched. Our knowledge of repolarization on contact comes from studies on flat surfaces, unlike cells in the body, which crawl along fibers. We discover that cells on single fibers walk past one another– but that cells in contact with multiple fibers stick to one another and move as pairs. This outcome changes to walk-past after cell division. Our experiments and models reveal how the environment regulates cell-cell coordination after contact.

## Introduction

Cell migration is an essential component of various physiological processes such as morphogenesis, wound healing, and metastasis^1^. Cell-cell interactions in which cell-cell contact reorients cell polarity are necessary for the correct function of many developmental events^2^. One of the earliest such interactions known was termed ‘contact inhibition of locomotion (CIL)’ by Abercombie and Heaysman over five decades ago in chick fibroblasts cultured on flat 2D substrates^2–4^. In CIL, two approaching cells isolated from the rest of cell population first make contact, followed by protrusion inhibition at the site of contact, which leads to cell repolarization through formation of new protrusions away from the site of contact. Subsequently, cells migrate away from each other in the direction of newly formed protrusions^1^. This sequence can however be altered in specific conditions such as metastasis where a loss of CIL allows malignant cells to invade fibroblast cultures – this is a loss of CIL between different cell types (heterotypic CIL)^4,5^. Recent work has also begun to identify the molecular players that initiate and regulate CIL, including Rac activity, microtubules, Eph/Ephrin binding and E- and N-cadherin expression^6–10^.

CIL is most commonly studied and analyzed on flat 2D substrates using several invasion and collision assays^2,3,11^. By contrast, cells traveling in matrix *in vivo* are constrained to move along narrow fibers. A common drawback in use of featureless 2D assays is thus the inability to study CIL under natural constraints^11–13^. Recently, micropatterned substrates have been used to understand restricted motility, developing 1D collision assays where cell migration is constrained to straight lines, allowing for a greater occurrence of cell-cell collisions to quantify rates and outcomes of different types of cell-cell interactions^11,13–15^. These interactions do not necessarily resemble the stereotyped CIL behavior. Broadly, experiments and simulations^16–18^ have observed 1) the classical stereotype of CIL with two cells contacting head-on, with both cells repolarizing (referred to as “reversal” or “mutual CIL”), 2) after a head-on collision, only one cell reverses (“training” or “non-mutual CIL”), and 3) cells manage to crawl past or over one another, exchanging positions (“walk-past” or “sliding”). Within the well-studied neural crest cell explants, walk-past is extremely rare^11^, but it can occur in epithelial cells, especially in those who have been metastatically transformed or with decreased E-cadherin expression^15^.

Both 2D substrates and micropatterned stripes provide controllable and reproducible environments, but neither fully models the details of *in vivo* native cellular environments, which consist of extracellular matrices of fibrous proteins, with these fibers having different radii. Our earlier *in vitro* recapitulation of the effects of fiber curvature showed that both protrusive and migratory behavior is sensitive to fiber diameter^19–21^. Furthermore, we have shown that suspended flat 2D ribbons do not capture the protrusive behavior observed on suspended round fibers^19^, thus, we wanted to inquire if the CIL rules developed on 1D collision and 2D assays extend to contextually relevant fibrous environments. To understand CIL in fibrous environments that mimic native ECM, we use suspended and aligned nanofiber networks to study CIL behavior in NIH/3T3 fibroblast cell-cell pairs exhibiting two distinct elongated morphologies, spindle, attached to a single fiber and parallel cuboidal, attached to two fibers^22^. We further investigate the effect of cell division on CIL by studying the encounters of cells that have recently divided (daughter cells) with other cells; these recently-divided cells are much faster, consistent with earlier work^23^. Our work allows us to determine the types and rates of cell-cell contact outcomes – the “rules of CIL” – in an experimentally relevant system with a controlled geometry. These rules are radically different from the known stereotypical behavior in 2D assays, but the essential features of these rules emerge robustly from a minimal computational model of CIL in confined geometries.

## Results

### Fiber spacing and diameter control cell shape; cell division controls speed

To develop the rules of CIL in fibrous environments mimicking native extracellular matrix (ECM), as shown schematically in **Figure 1a**^24^, we must first specify a controllable geometry. We cultured cells on suspended fibers of three diameters: 130, 500, and 1000 nm, which we created using the previously reported non-electrospinning spinneret based tunable engineered parameters (STEP) method (**Figure 1b**)^25–27^ We observed that cells on these fiber networks were generally found in a “spindle” geometry when placed on fibers with spacing ~20 μm where cells are only in contact with one fiber and elongate along that fiber, or in a “parallel cuboidal” geometry, spanning two fibers (**Figure 1c**), when on fibers with spacing ~10 μm. We selected ~500 nm diameter as a model system, as cells on ~130 nm exhibited dynamic plasticity in shape changes leading to shedding of cellular fragments from longer protrusions during migration (**Movie 1 and Figure 1c, S1**), and the protrusions formed on 1000 nm diameter fibers were of similar lengths as those observed on 500 nm diameter fibers but at times difficult to optically discern (**Movie 2**). We also immediately observed that cells underwent mitotic division subsequently had daughter cells with a large increase in speed in both spindle and parallel configurations (**Figure 1d**).

**Figure 1:**
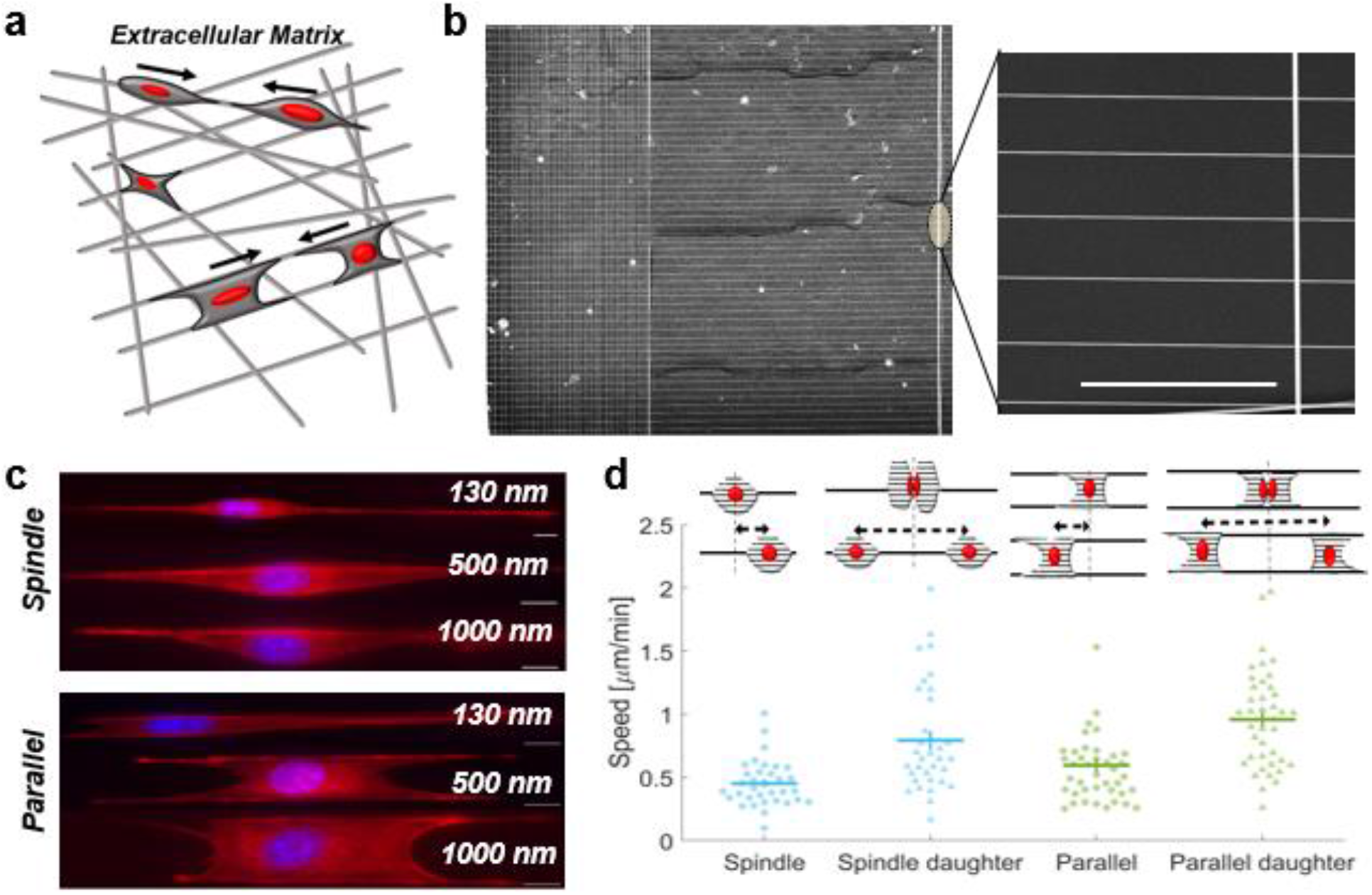
Cell motility on nanofiber network: **a)** Depiction of a cell’s *in vivo* extracellular matrix environment with ECM fibers. **b)** SEM image of aligned parallel fibers (500nm) manufactured using the STEP method and used for CIL experiments. Scale bar is 50μm. **c)** Spindle and parallel cuboidal cell morphologies on fibers of three diameters stained for actin (red) and nucleus (blue). Cells on 130 nm diameter fibers form long protrusions. **d)** Daughter cells have significantly increased speed compared to cells that haven’t recently divided (n=34 and 41 for each spindle and parallel case, respectively). Error bars show standard error.

### CIL collision outcomes in two approaching spindle cells

Next, we wanted to observe and quantify the outcomes of spindle cell CIL interactions. We looked at 47 randomly selected approaching spindle cell-cell collisions without cell division from 40 experiments. We found that spindle cells approaching each other mostly walked past each other without repolarizing (66%, **Figure 2a and Movie 3,4**). Since spindle cells on fibers form focal adhesions at the poles^22^, they can shift position around the fiber, thus allowing them to walk past one another. We also observed non-mutual CIL (30%) where upon contact only one spindle repolarized and both cells continued to migrate as a cohesive unit (“train”) in the new migration direction (**Movie 5**). Very rarely, 4% of the time, we see a mutual CIL response wherein upon contact, both spindle cells experience protrusion inhibition at the site of contact and repolarize away from one another – the stereotype of CIL on 2D substrates (**Movie 6**). In a separate set of collisions where one of the colliding cells had recently divided (n=98), we observed increased occurrence of walk-past behavior (82%, **Movie 7,8**) and a decrease in both non-mutual CIL (17%, **Movie 9,10**) and mutual CIL (1%). We also tracked the cell position over time for approaching spindles with and without cell division **(Figure 2b)**. The transient profiles reveal that in the scenario without cell division, there is no significant deviation in cell speed throughout the walk-past, though in the presence of cell division, the daughter cell slows down upon contact with the approaching spindle **(Figure 2c, d)**. Division increases the speed of daughter cells and also increases the rate of cells walking past each other – does the increased cell speed of daughter cells also lead to walk-past events that are shorter in duration? Surprisingly, we found almost no difference in the time in contact between spindle-spindle and spindle-daughter walk past events **(Figure 2e)**. Similarly, we do not find a large effect of division on the time for repolarization in non-mutual CIL (Figure S2).

**Figure 2:**
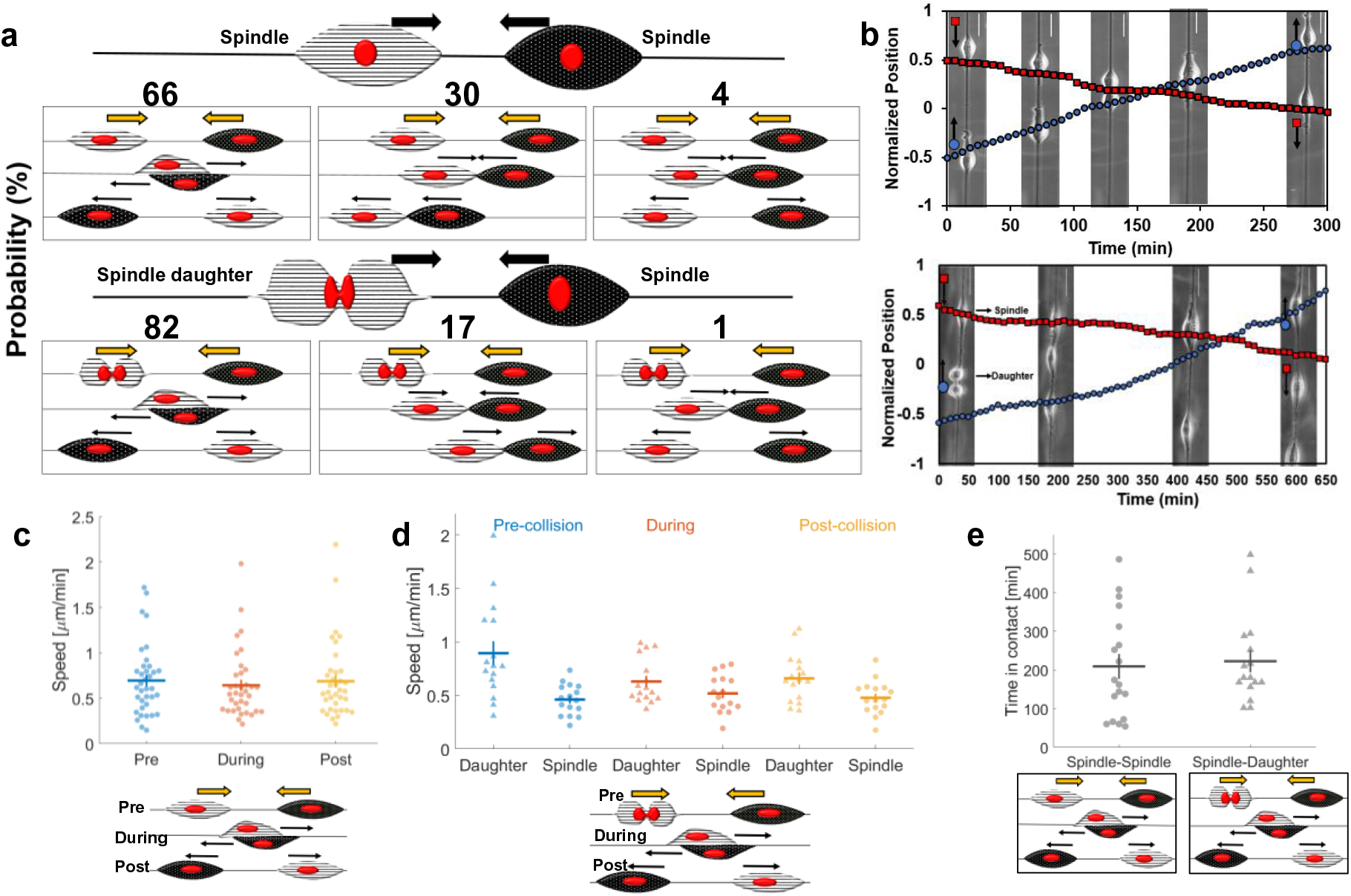
Spindle cell CIL: **a)** Outcomes of two approaching spindle cells in the absence (n = 47) and presence (n=98) of cell division. Walk-past is the dominant mode, followed by non-mutual (training) and mutual (reversal) CIL. **b)** Cell position and phase images over time for a walk past event without division (top) and with division (bottom). **c)** Average speed pre, during and post contact for spindle cells undergoing walk past (n=18 collisions). **d)** Average speed pre, during and post contact for spindle and daughter cells undergoing walk past (n=16 collisions). **e)** Graph comparing time in contact for spindle walk-past without and with cell division (n=18,16). Error bars show standard error.

### Parallel cuboidal cell CIL outcomes

Next, we wanted to explore the outcomes of two parallel cell collisions. We observed 28 randomly selected cell collisions without division from 41 independent experiments **(Figure 3a)**. In the absence of cell division, CIL outcomes were strikingly different than in the spindle geometry: two approaching parallel cells always resulted in non-mutual CIL (100% in 28 randomly selected movies, **Movie 11, 12**). One of the cells would repolarize and subsequently both cells would move in the new migration direction as a cohesive unit (training). In some instances, we observed that after the initial training phase, the repolarized cell would move faster leading to both cells separating but moving in same direction (**Movie 13**) and in rare instances one or both cells would repolarize (**Movie 14**). We have counted these outcomes as training as long as the initial motion lasted for at least one cell length. However, in the presence of cell division, from 59 randomly selected cell collisions, we observed two outcomes: walk-past (**Movie 15, 16)** and non-mutual CIL (**Movie 17**). The dominant outcome of collision between a faster moving daughter cell and a normal cell was walk-past (63%). During these walk-past events, we often observed that the daughter cell changed shape to a spindle cell (attached to one fiber) during the contact period. Post contact, the daughter regained its original parallel shape in most of cases observed. As done previously for spindle cells, we tracked the cell positions over time for both collisions involving recent post-division cells and those without (**Figure 3b**). Similarly, to our results in spindle cells, in the absence of cell division – when the only outcome is a train of cells being formed – cell speeds before and during contact were similar in both cells **(Figure 3c)**. Similar results were also seen in the presence of division **(Figure S3)**. We emphasize that, though cells can be briefly stalled during a collision (**Figure 3b**), the speed may not drop to zero, because speed measured as changes in both *x* and y centroids over a time of 6 minutes can reflect changes in cell shape as well as motility. Can variability in speed predict the outcome of repolarization? We found no evidence that the cell that repolarized was slower than the cell that did not **(Figure 3d)**. As in spindle cells, we found no strong dependence of the repolarization time of non-mutual CIL on the presence of cell division **(Figure S3)**.

**Figure 3:**
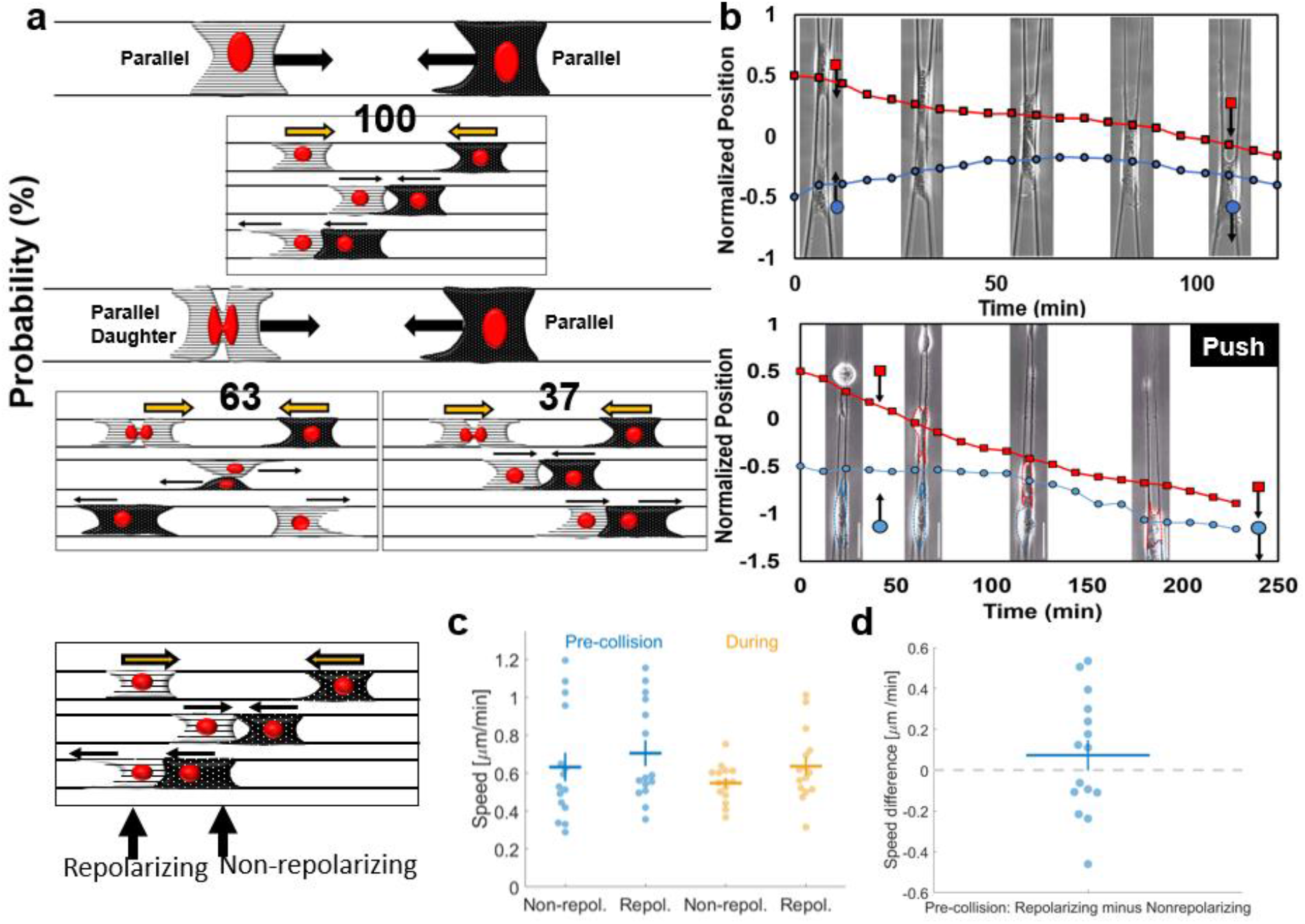
Parallel cell CIL: **a)** Outcomes of two approaching parallel cells in the absence and presence of cell division. Non-mutual CIL is the dominant mode in the absence of cell division – 100% of approaching parallel cells result in one cell repolarizing and altering its migration direction (n = 28). With cell division, walk past behavior is the most likely to occur followed by non-mutual CIL (n = 59). **b)** Cell position and phase images over time for parallel cells exhibiting non-mutual CIL (top) and ‘push’ (bottom). **c)** Average cell speeds pre and during contact for parallel non-mutual CIL for the cell that repolarizes and the non-repolarizing cell., **d)** No evidence that the cell that repolarizes has a slower or faster speed than the cell that does not. Error bars indicate the standard error.

In collisions with daughter cells, a rare subset of the non-mutual CIL had, unexpectedly, an apparent cell ‘*push*’, where a daughter cell that collides with a normal parallel cell leads to the non-daughter cell moving, even before the non-daughter parallel cell repolarizes and makes new protrusions **(Figure 3b, Movie 18, 19)**. This suggests that it is the physical force exerted by one cell literally pushing the other, overcoming any resistance from cell-fiber adhesion.

### Leading-Trailing spindle and parallel cell collisions

In addition to leading-leading or head-head collisions, we also observed instances where the leading edge of one cell would contact the trailing edge of another cell and both cells would continue to migrate (spindle-spindle and parallel-parallel). This behavior was previously reported in collisions on micropatterns^13^ and suggests that there may be an asymmetry in the signals cells receive in contact with another cell’s tail or head. Recent experiments describing “contact following of locomotion” also support the idea of an asymmetry of interactions^28^.

We show the speeds of the trailing and leading cells prior to the collision and then during the contact in **Figure 4a** for all of the collision types (spindle-spindle: **Movie 20**, spindle-daughter: **Movie 21**, parallel-parallel: **Movie 22**, and parallel-daughter: **Movie 23**). Due to the variability of cell speeds, some information is not apparent in this data, but is only clear in paired comparisons. For instance, we show in **Figure 4b** that the trailing cell always has higher speed than the leading cell – which is necessary for the trailing cell to catch up. Interestingly, the primary effect of a trailing cell catching up with a leading cell is that the leading cell gains speed during contact **(Figure 4c)**. This effect is consistent over all of the types of collisions we studied.

**Figure 4:**
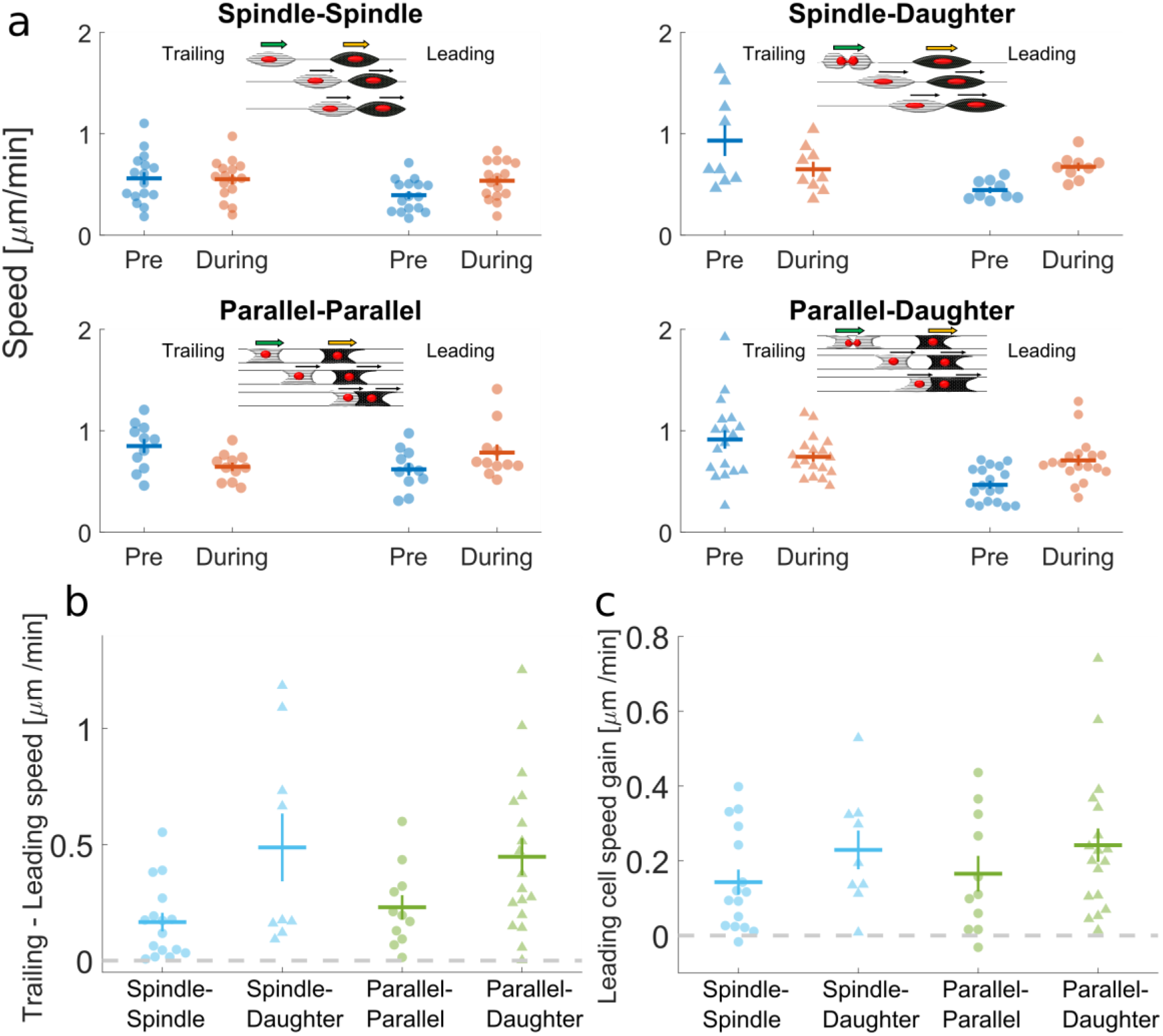
Leading-trailing CIL: **a)** Average speeds for cells before and during a trailing-leading contact. Four cases are shown: i) spindle-spindle collisions, ii) spindle-daughter spindle collisions, iii) parallel-parallel collisions, and iv) parallel-daughter parallel collisions. **b)** Speeds of trailing cells are always larger than that of leading cells, allowing them to catch up. **c)** Leading cells robustly gain speed once they are in contact with the faster trailing cell.

### A simple simulation framework captures CIL outcomes

Our experimental results show three broad qualitative features which differ from classical results in two-dimensional CIL: 1) spindle-spindle collisions commonly lead to walk-past; 2) the parallel fiber geometry suppresses this walk-past behavior, and 3) cells that have recently divided, which move more quickly, are more likely to walk past another cell – even within the parallel geometry. Altering the adhesive environment of a cell can regulate many processes^29^, while cell division also can alter the localization of receptors on the cell surface and ability to respond to signals^30^, motility and the extent of contact guidance^23^. We want to understand to what extent our observed CIL features can arise *solely* from the change in geometry and the observed change in speed of recently-divided cells. To address this question, we developed a theoretical framework for our fiber CIL by extending our previously reported 2D CIL model^31^. This model describes cells with positions **r**_i_ and polarities **p**_i_. The polarity is a vector indicating the direction a cell is traveling – its magnitude is the speed the cell would have in the absence of other forces acting on it. Cells are modeled as self-propelled particles, with forces attaching them to each of the fibers. Between the two cells are adhesive forces representing cadherin-mediated attachment, and forces of repulsion to represent elastic deformation of cells when they begin to contact. In addition, we include two key biochemical processes affecting the cell’s polarity: 1) contact guidance, in that cells tend to polarize along the fibers, and 2) contact inhibition of locomotion, in which cells tend to polarize away from contact with the fronts of other cells. In addition, we include stochastic fluctuations in polarity. Cell division is modeled solely by increasing the initial polarization (speed) of the cell. The full model equations are shown in the Materials and Methods below, with additional details and parameters in the Supplementary Information.

We used our model to simulate cell-cell collisions (**Figure 5a**), and characterized three outcomes: 1) walk-past, where cells crawled past each other, exchanging positions, and then separating, 2) reversal, where cells contacted each other, repolarized, and separated without exchanging positions (“mutual CIL”, and 3) training – where the two cells failed to separate, and generally had one repolarizing cell following the other (“non-mutual CIL”).

**Figure 5:**
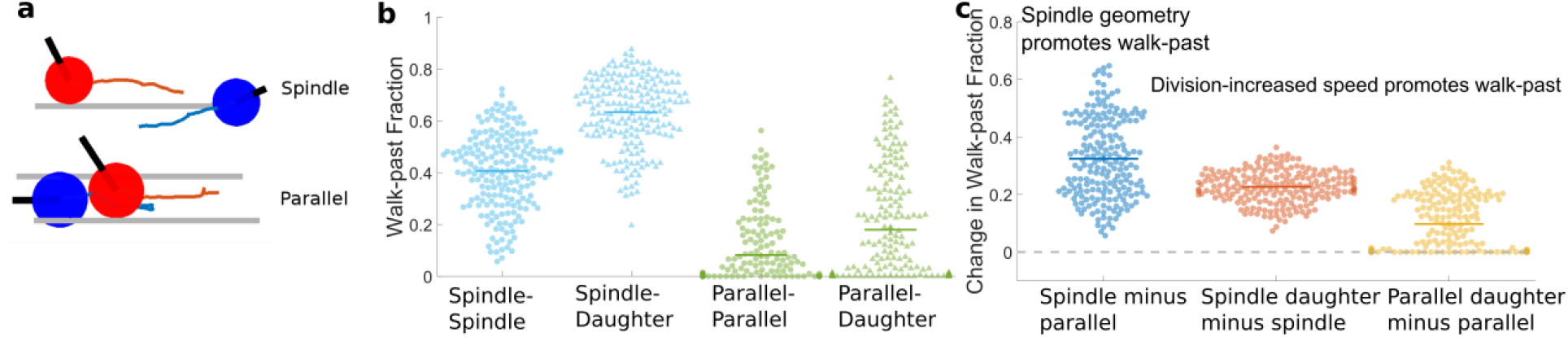
Model reproduces key qualitative experimental outcomes robustly to parameters: **a)** Illustration of model for cells on single fiber (spindle geometry) and attached to two fibers (parallel geometry). Black lines indicate cell polarity **p**; grey lines indicate fibers. The red and blue lines indicate the past trajectory of the cells. A walk-past event is shown for the spindle configuration, and a training event is shown for the parallel configuration. **b)** The fraction of walk-past events is shown for parameters randomly sampled from plausible values; each dot is a different parameter set. Though changing parameters affects the overall fraction of walk-past, walk-past is common. **c)** Comparisons between walk-past levels. Holding other parameters fixed, the spindle geometry always has more walk-past than the parallel. Collisions where division occurs always have an increase in walk-past for spindle, and never have a decrease in walk-past behavior in parallel geometry.

We find that, in the spindle geometry, simulated cells can walk past each other (**Figure 5b, Movie 24, 25**). In large part, this reflects that cells can freely rotate around the fiber, and that they can therefore either rotate after contact or, if the cells start at different angles, only come in glancing contact. The details of the cell-cell contact and possible rotation are difficult to distinguish in microscopy, but from videos it is clear that cells can have their bodies distributed to one side of the fiber and can shift their body to other side during migration. When simulated cells are attached to two fibers, cell body distribution around the sides of the fiber is strongly suppressed, and walk-past occurs less frequently (**Figure 5c, Movie 26, 27**) This behavior is generic, and is true across many parameters; we show in Figure 5c that, sweeping across plausible parameters, the fraction of cells that walk past one another is always lower in the parallel geometry than in the spindle geometry for 200 independent parameter sets spanning the plausible range of values. Similarly, we see that when one cell has recently divided (modeled as having a higher initial velocity – multiplying its polarization in the x direction by a factor of three; this is roughly set by the increase in mean velocity in the x direction observed experimentally), walk-past is almost always increased in the spindle case, and never decreases in the parallel case. Some parameter sets for parallel geometries with very high attachment to the fiber have zero change in walk-past rate due to division, because walk-past is effectively impossible due to the tight constraints – this is the excess of zeros in the third column of Figure 5c. We thus find that, at a qualitative level, matching behavior of cell-cell collision outcomes seen in our experiments and simulations are robust, showing that geometric confinement alone is sufficient to alter outcomes of CIL. In addition, our simulations show that *in silico*, it is sufficient to merely increase a cell’s speed in order to increase walk-past.

Our central results are robust to many model parameters, but the quantitative values vary with the specific parameters chosen (**Figure 5**). We fit our model to the experiment by varying the five parameters where we do not have pre-existing values (strength of CIL *β*, cell-cell adhesion strength *α*, cell-fiber attachment strength *κ*, cell stiffness *κ_cell_*, and variation in cell velocity *σ*) over plausible ranges, and choose the best fit. Nonetheless, our model is in excellent agreement with experimental observations of spindle head-head collisions, and reasonably agrees well with the parallel fiber category (**Figure 6**), thus allowing us to qualitatively describe rules in CIL outcomes on suspended fibers.

**Figure 6:**
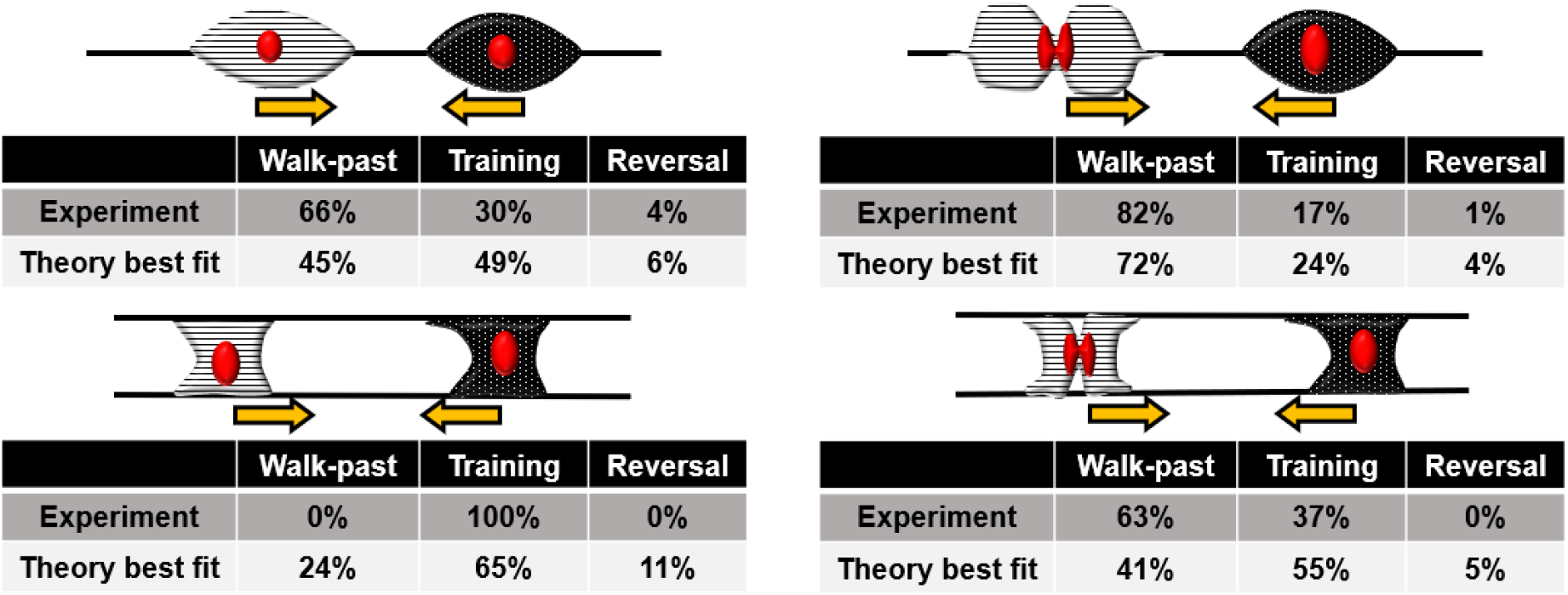
Summary of comparison between experiment and best-fit theory. Theory values are for the best-fit parameters provided in the Supplemental Information and are generated from statistics of N = 500 collisions.

## Discussion

In this paper, we present a combined experimental-theoretical platform to quantify outcomes of cell-cell collisions and CIL in fibers analogous to *in vivo* extracellular matrix. Using suspended fibers in aligned patterns, we investigated CIL decisions of two fibroblasts approaching each other in two elongated shapes: spindle cells attached to single fibers, and parallel cuboidal cells attached to two fibers. **(Figure 1)**. We observe that two approaching spindle cells most commonly do not repolarize but rather walk past one another and continue along their respective migration direction **(Figure 2)**. We attribute this behavior to spindle cells being able to shift their position about the fiber axis, as they are primarily attached to the fibers at the poles, as demonstrated by a gentle push to a spindle cell using an external probe that swivels the spindle cell about the fiber axis (**Movie 28**). Spindle walk past behavior is similar to a rare walk-past observed in neural crest cells on micropatterned lines where the cells also pass by one another without repolarization^11^, though notably, this walk past behavior was extremely rare (under 2% of collisions of wildtype cells, and about 10% when CIL was suppressed). Milano et al. have also observed a similar “sliding” behavior – though they find that it is only present at high rates (~50%) in metastatic MDA-MB-231 cells, and the rates of sliding can be increased by downregulation of E-cadherin in noncancerous MCF-10A. By contrast to both of these results, our experiments show that a walk-past behavior can be the majority outcome even in normal 3T3 cells on suspended fibers. We can also regulate this behavior by changing the local geometry: when two parallel cells approach one another, upon contact, one cell will repolarize, switch its migration direction and continue to move as a unit with the other – non-mutual CIL or train formation **(Figure 3)**. These behaviors differ from the traditional CIL definition as both cells do not inhibit their protrusive behavior and alter their migration path as seen in initial studies^4^, though similar behavior has been captured before both *in vitro^13^* and *in vivo^32^.* Our simulations (**Figures 5–6**) also suggest that the transition from a high rate of walk-past to a high rate of train formation can be generated solely by changing the geometry of the extracellular matrix from single to two parallel fibers. Importantly, we hold all other parameters in the model constant – we do not assume that the changed matrix geometry alters CIL strength explicitly. Cell division has significant influences on both spindle and parallel CIL. Both spindle and parallel daughter cells migrate at a significantly faster speed post division than that of a non-daughter cell, and when a daughter cell comes in contact with another cell, walk-past occurs more often – in both parallel and spindle geometries. Interestingly, in the parallel geometry, this walk-past generally occurs with a parallel cell temporarily detaching from one fiber, and is thus able to go around the parallel cell in spindle configuration. Our simulations suggest that the increased speed of daughter cells alone may be sufficient to explain the increased rate of walk-past – within our model, walk-past rates robustly increase when one cell’s initial polarization (speed) is increased. However, we should note that cell division will likely also affect the degree of cell adhesion, and that prior work has emphasized that cues from mitosis can override chemical signals, suggesting that division may have independent effects on CIL^30^.

We also note an important effect of head-tail collisions of cells in changing cell speed, in which a trailing cell catches up with a leading cell. When a trailing cell – which must be moving faster than the leading cell to catch it – contacts the leading cell, the leading cell almost always speeds up (**Figure 4c**). This effect would be anticipated from simple models of cells as self-propelled particles, which suggest that the velocity of a cluster of cells is the average of the velocity they would have if they were separated^31^.

One rare but intriguing phenomenon we observe in parallel cell CIL is that of a cell “push.” Here, two parallel cells approach one another, and one cell will undergo division. Following division, when a daughter and parallel cell contact we have seen non-mutual CIL event in which the non-daughter cell moves even before its protrusions are repolarized away from contact **(Figure 3)**. This shows that – in extreme cases – a daughter cell might be able to physically push another cell, overcoming the cell-fiber adhesion strength.

While our model, robustly to parameters, reproduces the key qualitative outcomes (**Figure 5**), our best fit to the observed quantitative rates does not perfectly reproduce the experimental outcomes (**Figure 6**). We emphasize that, because we have not modeled the effect of cell shape in our simulations, and that we have not addressed cell-to-cell variability, which we expect to be relevant in these events^17^, we should not expect a perfect result. Nevertheless, we have obtained a reasonable best fit. The most challenging outcomes to predict with a single consistent set of parameters are the experimental observations that 0% of cells walk past in the parallel geometry without division while 63% of them walk past when division occurs. It is possible to choose parameters so that the cells are so tightly adherent to the fibers that they cannot walk past one another (**Figure 5**), but when this is done, the increase in speed alone is insufficient to create walk-past in the parallel geometry (**Figure 5c**). We suspect that this difficulty in fitting largely reflects our assumption that cells remain connected to both fibers, whereas in experimental observations, we observe that parallel walk-past often occurs with one cell becoming disconnected from one fiber, and becoming more spindle-like. To provide more insight into the roles of different parameters in the model, we take these fitting parameters as our “wild type” and show how the outcomes depend on key parameters like cell-cell adhesion (**Supplementary Information**).

In conclusion, in this paper, we present a platform to demonstrate how CIL depends on extracellular matrix geometry and cell geometry in a system closely analogous to extracellular matrix fibers. Our results are strikingly different from prior studies on CIL, observing cells able to walk past one another even in healthy, non-invasive cells. We show, using computational modeling, that this can be explained solely as a function of the different fiber configuration we expose cells to, suggesting that simple micropatterned lines may not be sufficient to capture *in vivo* cell-cell interactions in extracellular matrix. This has implications beyond our initial focus on homotypic interactions between two fibroblast cells. Loss of heterotypic CIL is viewed as a signature of cancerous cells, allowing cancerous cells to invade the healthy population^14,33^; neural crest cells also maintain homotypic CIL among themselves while lacking a heterotypic CIL response allowing them to invade the mesoderm layer and other tissues^1^. Our results imply that earlier CIL findings from use of flat 2D substrates, microprinted lines, and confined channels, for both homotypic and heterotypic cell-cell interactions, may need to be re-examined in order to better understand the role of extracellular matrix geometry.

## Materials and methods

### Nanofiber network manufacturing

A 10% wt. solution was made by dissolving polystyrene ((PS) MW: 2,000,000 g·mol^-1^) in xylene. Migration networks of nanofibers with ~500nm diameters with 15μm or 30μm spacing were created using the non-electrospinning Spinneret based Tunable Engineered Parameters (STEP) method. After formation, the nanofibers were fused by exposure to tetrahydrofuran (THF) utilizing a custom fusing chamber. Fibers were deposited at two spacing: ~ 20 μm to achieve spindle cells that attached to single fibers, and ~10 μm to achieve parallel cuboidal cells attached to two fibers.

### Cell seeding and imaging

The nanofiber networks were placed in individual wells of a six-well plate and sterilized before coating with 4μg/mL of fibronectin for approximately one hour. NIH3T3 fibroblasts were cultured in Dulbecco’s Modified Eagle Medium (DMEM) supplemented with 10% Fetal Calf Serum (FCS) at 37°C and 5% CO2 in T-25 flasks. The cells were seeded at a low concentration onto the nanofiber networks after removal of the fibronectin and were subsequently left to attach for one hour prior to flooding each well with 3mL of media. Time-lapse imaging for migration was executed for 12-24 hours with an imaging interval ranging from 1 minute, 2 minutes or 3 minutes using a 10x or 20x objective.

### Cell migration analysis

Time-lapse videos were created with spindle and parallel cuboidal cells prior to contact, during contact, and for some cases post contact. Image J was used to outline cells, track cell centroids every six minutes and obtain an average migration rate (μm/min). The x and y coordinates of cell centroid were measured at each time point and the 2D speed was determined by (Vx^2^+Vy^2^)^1/2^. Normalized cell position vs. time transient plots were created by measuring the cell positions relative to the initial midpoint of the two cells being analyzed, rescaled by the starting distance between their centroids. Only clean cell-cell collisions (those where only two cells interacted) were used for analysis in order to avoid multiple cell interferences.

### Statistical analysis

Throughout the paper, we show error bars corresponding to the standard error (standard deviation of the mean).

### Computational model

Cells are characterized by a position **r**_i_ and a polarity vector **p**_i_. The vector **p**_i_ reflects the degree of biochemical polarization of cell *i* – this indicates both the orientation of the cell’s motion and its speed – more polarized cells have larger velocity. In the absence of additional forces on the cell, cell *i* travels with velocity **p**_i_:

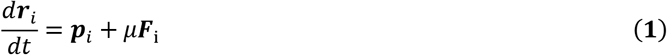

Here, **F**_i_ is the force exerted on cell *i* by other cells and the environment, and *μ* is the cell’s mobility, which relates forces F exerted onto the cell to the resulting velocity. The polarity evolves by the stochastic equation:

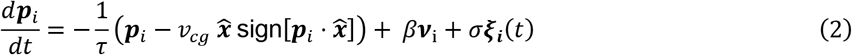

The first term in this equation states that polarity relaxes to its equilibrium value over a timescale τ, and that its equilibrium value – in the absence of other cells – is 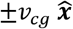, i.e. the cell will be traveling along the x direction (fiber direction) with speed *ν_cg_*. This approach lets us model contact guidance: cells will tend to polarize along the fiber direction. The term sign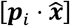 ensures that cells that are weakly polarized along +/- x become more strongly polarized in this direction. The second term *β**ν**_i_* models contact inhibition of locomotion, biasing the polarization of each cell away from its neighbors. However, unlike our earlier model, we choose to treat CIL as asymmetric – contact with a neighboring cell’s front (leading edge) repolarizes the cell, but contact with a neighboring cell’s rear (trailing edge) does not. This is represented as:

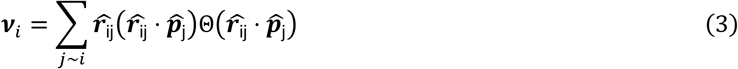

where the sum is over cells *j* contacting cell *i* (*j ~ i* meaning those within radius *R* of cell *i*), 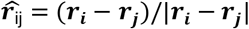 is the unit vector connecting cells *i* and *j*, and 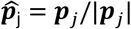. Here, Θ(*x*) is the Heaviside step function, which is zero for x<0 and one for x>0. With this model, ***ν***_i_ is a vector that points away from the cells whose fronts contact cell *i*, weighted more toward those who are directly pointing at cell *i*. This asymmetry between front and rear contact has been experimentally observed^13,28^, and is consistent with our experimental observation of train formation. However, we note that symmetric interactions can also lead to train formation^34^. The final term in Eq. (2) is a stochastic Gaussian Langevin noise modeling random changes in polarization.

To complete the model, we need to specify the cell-cell forces and the cell-environment forces **F**_i_, which are found as the derivative of a potential,

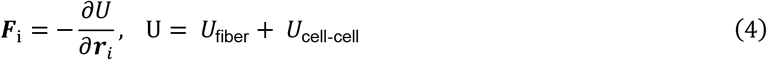

We model an adhesion force between the cell and the fiber using a harmonic force that prefers to keep the cell’s center at a distance R from each fiber (R here being a cell radius):

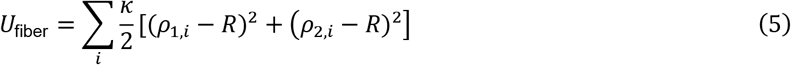

Where 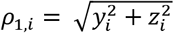 is the distance from the ith cell to the first fiber and 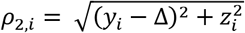 is the distance from the ith cell to the second fiber – Δ is the fiber separation. This is chosen as a minimal option making cells adherent to the fiber, and is parameterized by the spring stiffness *κ*. (For spindle cells, only the first term in Eq. 5 is kept.)

Cell-cell interactions *U*_cell-cell_ are composed of a short-range repulsion, representing the elastic deformation of the cells as they become closer to one another, and a longer-range adhesion, which states that for cells in contact to be pulled apart, force must be applied to them. These potentials are:

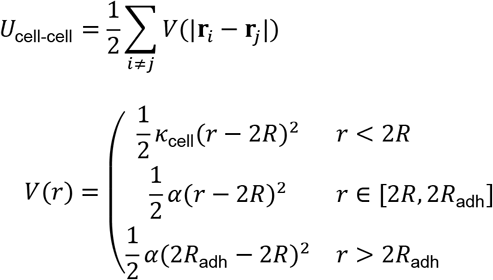

where *κ*_cell_ is the spring constant of the cell-cell interaction, *α* characterizes the strength of adhesion, and *R*_adh_ is the range of adhesion.

To investigate the statistics of front-front collisions, we first initialize the model by simulating single cells, turning off all cell-cell interactions. We then keep the cell polarities fixed, but place the cells so that they are facing each other and their centers are separated by three cell radii, turn on cell-cell interactions, and restart the simulation. We ran the simulation multiple times and tracked the statistics of different outcomes.

Additional computational details, including numerical values of parameters, are available in the Supplementary Information.

## Acknowledgements

ASN would like to acknowledge the Institute of Critical Technologies and Sciences (ICTAS) and Macromolecules Innovation Institute (MII) at Virginia Tech for their support in conducting this study. ASN would like to thank STEP Lab members and undergraduate student Megan Dobbins for their helpful suggestions and discussions. This work is supported by National Science Foundation (1762634) awarded to ASN. BAC acknowledges support from Johns Hopkins University and the National Science Foundation (PHY 1915491).

## Author contributions

ASN conceived the study. BC and ASN designed the study. JS and ASN conducted experiments. JS, BC and ASN analyzed data. BC developed and implemented the theoretical model. JS wrote the manuscript. BC and ASN edited and finalized the manuscript.

## Movie Legends

Movie 1: Protrusions on suspended small (135 nm diameter) fibers. Cells form long protrusions on suspended small diameter fibers. Long protrusions often break from cell body resulting in fragments (white arrows) attached to fibers. Time shown in hours:minutes:seconds:thousandths

Movie 2: Protrusions formed on suspended large (1000 nm diameter) fibers. Protrusions formed from cells attached to suspended large diameter fibers can be difficult to discern optically. Time shown in hours:minutes:seconds:thousandths

Movie 3,4: Walk past of spindle shaped cells on suspended fibers (500 nm diameter). Two spindle shaped cells attached to suspended fibers approach each other, make contact and walk past. Time shown in hours:minutes:seconds:thousandths

Movie 5: Non-mutual CIL in spindle cell pairs on suspended fibers (500 nm diameter). Two spindle shaped cells attached to suspended fibers approach each other and following contact the cell on left (black arrow) repolarizes. Subsequently both cells move together resulting in non-mutual CIL. Time shown in hours:minutes:seconds:thousandths

Movie 6: Mutual CIL in spindle cell pairs on suspended fibers (500 nm diameter). Spindle shaped cells attached on suspended fibers approach each other and following contact both cells repolarize resulting in mutual CIL. Time shown in hours:minutes:seconds:thousandths

Movie 7,8: Walk past of spindle shaped cell and daughter cell post division on suspended fibers (500 nm diameter). A spindle shaped cell and a fast-moving daughter spindle shaped cell on suspended fibers approach each other, make contact and walk past. Time shown in hours:minutes

Movie 9. Non-mutual CIL of spindle shaped cell and daughter cell post division on suspended fibers (500 nm diameter). A slow-moving spindle shaped cell (white arrow) and a fast-moving daughter spindle shaped cell (black arrow) on suspended fibers approach each other, and following contact, the non-dividing cell on right repolarizes. Subsequently both cells move together resulting in non-mutual CIL. Time shown in hours:minutes:seconds:thousandths

Movie 10. Non-mutual CIL of spindle cell and daughter cell post division on suspended fibers (500 nm diameter). A slow-moving spindle cell (white arrow) and a fast-moving daughter spindle cell (black arrow) approach each other, and following contact, the daughter cell on left repolarizes. Subsequently both cells move together resulting in non-mutual CIL. Time shown in hours:minutes:seconds:thousandths

Movie 11,12. Non-mutual CIL of parallel-parallel cell pairs on suspended fibers (500 nm diameter). Two parallel cells on suspended fibers approach each other followed by one of the cells (black arrow) repolarizing. Subsequently both cells move together as a train for at least one cell length resulting in non-mutual CIL. Time shown in movie 11 in hours:minutes:seconds:thousandths and in 12: hours:minutes

Movie 13. Non-mutual CIL of parallel-parallel cell pairs on suspended fibers (500 nm diameter). Two parallel cuboidal shaped cells on suspended fibers approach each other followed by one of the cells (black arrow) repolarizing. Subsequently both cells move together as a train for at least one cell length followed by one of the cells (black arrow) separating. Time shown in hours:minutes:seconds:thousandths

Movie 14. Non-mutual CIL of parallel-parallel cell pairs on suspended fibers (500 nm diameter). Two parallel cuboidal shaped cells attached on suspended fibers approach each other followed by one of the cells (black arrow) repolarizing. Subsequently both cells move together as a train for at least one cell length followed by one of the cells (white arrow) repolarizing. This results in both cells moving apart from each other. Time shown in movie in minutes

Movie 15. Walk past of parallel cell in parallel-parallel cell pairs with one cell undergoing division on suspended fibers (500 nm diameter). One parallel cuboidal shaped cell on suspended fibers divides and the daughter cell approaches another parallel cuboidal shaped cell. The daughter cell changes shape to become a spindle shaped cell to achieve walk past. Time shown in hours:minutes:seconds:thousandths

Movie 16: Walk past of parallel cell in parallel-parallel cell pairs undergoing division on suspended fibers (500 nm diameter). Two parallel cuboidal shaped cells attached on suspended fibers divide and their respective daughter cells approach each other followed by both daughters changing to spindle shape to achieve walk past. Time shown in hours:minutes:seconds:thousandths

Movie 17. Non-mutual CIL of parallel cell in parallel-parallel cell pairs undergoing division on suspended fibers (500 nm diameter). Two parallel cuboidal shaped cells attached on suspended fibers divide and their respective daughter cells approach each other followed by one of the daughters (black arrow) repolarizing resulting in non-mutual CIL with training. Time shown in hours:minutes:seconds:thousandths

Movie 18, 19: Push of a parallel by a daughter parallel cell on suspended fibers (500 nm diameter). A dividing daughter cell (white arrow) of a parallel cuboidal cell moving at high speed physically pushes another parallel cuboidal shaped cell (black arrow). Push can be seen by movement of the pushed cell body before formation of visible protrusions. Time shown in hours:minutes:seconds:thousandths

Movie 20: Leading-trailing edge interactions between spindle cells on suspended fibers (500 nm diameter). A faster moving spindle shaped cell (black arrow) catches up and contacts the trailing edge of slower moving spindle (white arrow) shaped cell resulting in an increase in speed of slower cell. Time shown in hours:minutes:seconds:thousandths

Movie 21: Leading-trailing edge interactions between spindle cells on suspended fibers (500 nm diameter). A faster moving daughter spindle shaped cell (black arrow) catches up and contacts the trailing edge of slower moving spindle (white arrow) shaped cell resulting in an increase in speed of slower cell. Time shown in hours:minutes:seconds:thousandths

Movie 22: Leading-trailing edge interactions between parallel cells on suspended fibers (500 nm diameter). A faster moving parallel cuboidal shaped cell (black arrow) catches up and contacts the trailing edge of slower moving parallel (white arrow) cuboidal shaped cell resulting in an increase in speed of slower cell. Time shown in hours:minutes:seconds:thousandths

Movie 23: Leading-trailing edge interactions between parallel cells on suspended fibers (500 nm diameter). A faster moving daughter parallel cuboidal shaped cell (black arrow) catches up and contacts the trailing edge of slower moving parallel (white arrow) cuboidal shaped cell resulting in an increase in speed of slower cell. Time shown in hours:minutes:seconds:thousandths

Movie 24. Simulation of spindle-spindle collision showing walk-past. Cell polarity vectors are shown as black lines, and the red, blue cell trajectories over time are traced with the matching color. The fiber is shown as a gray line.

Movie 25. Simulation of spindle-daughter spindle collision showing walk-past. Cell polarity vectors are shown as black lines, and the red, blue cell trajectories over time are traced with the matching color. The fiber is shown as a gray line.

Movie 26. Simulation of parallel-parallel collision showing training. Cell polarity vectors are shown as black lines, and the red, blue cell trajectories over time are traced with the matching color. The fiber is shown as a gray line.

Movie 27. Simulation of parallel-parallel daughter collision showing walk-past. Cell polarity vectors are shown as black lines, and the red, blue cell trajectories over time are traced with the matching color. The fiber is shown as a gray line.

Movie 28. External probe pushing on a spindle cell on a suspended fiber (500 nm diameter). A computer-controlled glass probe pushes on a spindle shaped cell causing it to move about the suspended fiber

