## Supplementary Appendix for "Rules of Contact Inhibition of Locomotion for Cells on Suspended Nanofibers"

### I. SUPPLEMENTARY FIGURES

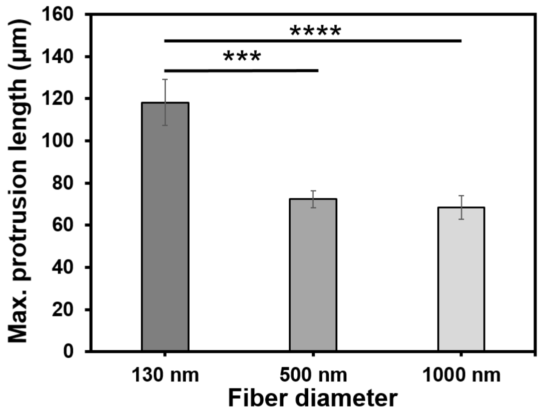


FIG. S1. Maximum protrusion length of cells on fibers of diameter 130 nm, 500nm, and 1000nm. (****, $p < 0.0001$; ***, $p < 0.001$; **, $p < 0.01$; *, $p < 0.05$).

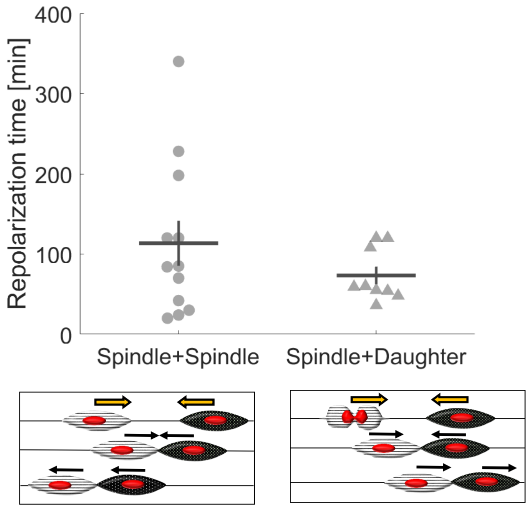


FIG. S2. Time to repolarize for spindle-spindle and spindle-daughter collisions that lead to train formation. $n = 12, 9$ respectively.

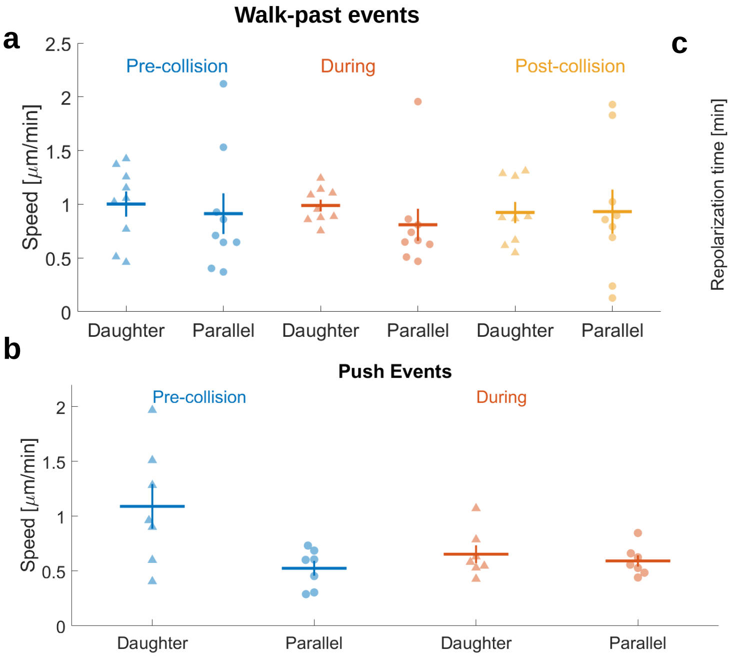


FIG. S3. Speeds and repolarization times for parallel collisions. **a**) Speed during walk-past events, which necessarily require one cell to be a daughter. **b**) Speed during "push" events where trains are formed but one cell is not immediately seen to repolarize. **c**) Comparison between repolarization time for parallel-parallel parallel-daughter, and parallel-daughter push events.

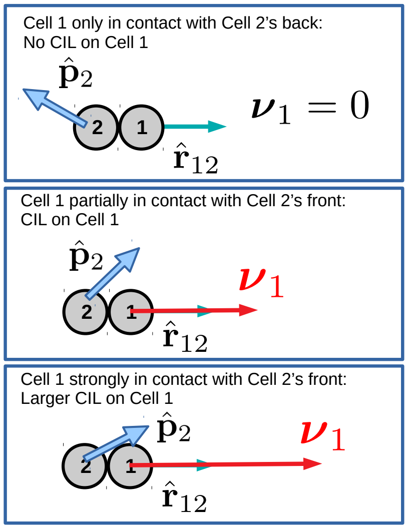


FIG. S4. Relative strengths of CIL in model depending on orientation of cells. We note that generally, cells are oriented along the $\pm x$ direction, so the orientations shown are not particularly realistic.

### II. ADDITIONAL MODEL DETAILS

We extend the model of [1] to describe interacting cells attached to a fiber. We model cells $i = 1, 2$ as having a polarity $\mathbf{p}_i$. Polarity is the velocity a cell *would* travel with if there were no mechanical forces acting on the cell (e.g. from other cells, the fiber) – thus its units are of velocity. This polarity obeys an equation:

$$\frac{d}{dt}\mathbf{p}_i = \underbrace{-\frac{1}{\tau}\mathbf{p}_i}_{\text{Relaxation}} + \underbrace{\frac{1}{\tau}\hat{\mathbf{x}}v_{cg}\frac{\mathbf{p}_i \cdot \hat{\mathbf{x}}}{|\mathbf{p}_i \cdot \hat{\mathbf{x}}|}}_{\text{contact guidance}} + \underbrace{\beta\boldsymbol{\nu}_i}_{\text{CIL}} + \underbrace{\sigma\boldsymbol{\xi}_i(t)}_{\text{Gaussian noise}} \tag{S1}$$

There are four terms:

- **Relaxation.** In the absence of any other terms, polarization relaxes to zero with the time $\tau$. This provides a characterization of the cell's persistence time in one direction.

- **Contact guidance.** Promotes polarization along the fiber axis (which we choose as $\hat{\mathbf{x}}$). If $p_x > 0$, this term polarizes the cell along $+\hat{\mathbf{x}}$, otherwise along $-\hat{\mathbf{x}}$. The term $\tau^{-1}$ in front means that, with only the first two terms, the cell will tend to relax to $p_x = \pm v_{cg}$.

- **CIL.** This term biases cells away from contact with their neighbors, into the direction $\boldsymbol{\nu}_i$:

$$\boldsymbol{\nu}_i = \sum_{j \sim i} \hat{r}_{ij} \left(\hat{r}_{ij} \cdot \hat{\mathbf{p}}_j\right) \Theta\left(\hat{\mathbf{r}}_{ij} \cdot \hat{\mathbf{p}}_j\right) \tag{S2}$$

  where the sum is over cells $j$ neighboring $i$ ($j \sim i$), i.e. those within a distance $2R$ from the cell, $\hat{r}_{ij} = (\mathbf{r}_i - \mathbf{r}_j)/|\mathbf{r}_i - \mathbf{r}_j|$ is the unit vector connecting cells $i$ and $j$ and $\hat{\mathbf{p}}_i = \mathbf{p}_i/|\mathbf{p}_i|$. $\Theta(x)$ is the Heaviside step function, $\Theta(x) = 0$ for $x < 0$, $\Theta(x) = 1$ for $x > 0$. The intuition for this term is shown in Figure S4 – the direction $\hat{\mathbf{r}}_{ij}$ points away from the neighbor $j$, modeling CIL – but CIL only happens when a cell is in contact with the front of another cell. The term $\hat{\mathbf{r}}_{ij} \cdot \hat{\mathbf{p}}_j$ ensures that this CIL will be stronger if the front is more aligned with the cell-cell contact axis – representing that CIL will be stronger if more of the front of one cell is in contact with one another. **This term differs from that in [1] because we assume only contact with another cell's front leads to repolarization.** This was found to be important in more complex models like [2, 3] and is one way for a CIL-like mechanism to create streams of cells.

- **Gaussian noise**. This last term is a white Gaussian Langevin noise; this fluctuation would drive the cell to have a persistent random walk in the absence of other features. With just relaxation and noise, the resulting "Ornstein-Uhlenbeck" model [4, 5] would have a Gaussian distribution of $\mathbf{p}$.

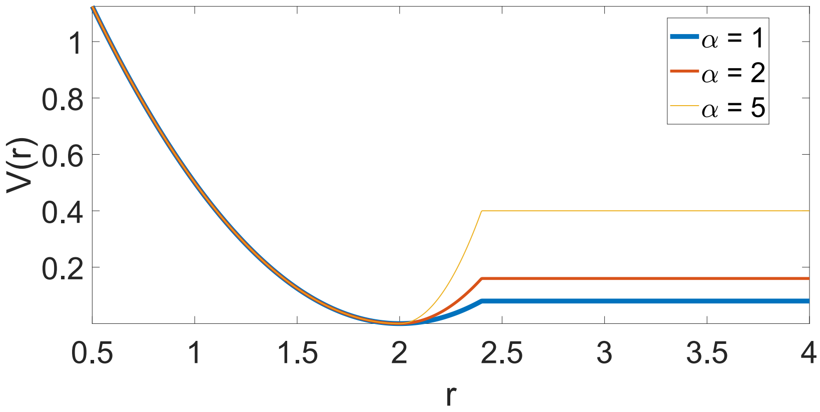


FIG. S5. Cell-cell interaction potential includes short-range repulsion and short-range attraction. Beyond the distance $R_{\text{adh}}$, the potential is constant (zero force). This is only illustrative and does not match the parameters in the paper – this graph shows $R = 1$ and $R_{\text{adh}} = 1.2$.

How does the polarity $\mathbf{p}_i$ of each cell translate into its motion? The motion of cells is at low Reynolds number (highly overdamped) so a force on the cell is translated into a constant velocity. We then write the cell's velocity as

$$\frac{d}{dt}\mathbf{r}_i = \mathbf{p}_i - \mu\frac{\partial U_{\text{fiber}}(\mathbf{r}_i)}{\partial \mathbf{r}_i} - \mu\frac{\partial U_{\text{cell-cell}}(\{\mathbf{r}\})}{\partial \mathbf{r}_i} \tag{S3}$$

where $\mu$ is a mobility coefficient for the cell, i.e. a constant force $F_z$ on a cell leads to a constant velocity $v_z = \mu F_z$. The term $U_{\text{fiber}}$ keeps the cell pinned to the fiber(s):

$$U_{\text{fiber}}(\mathbf{r}) = \frac{\kappa}{2}\left[(\rho_1 - R)^2 + (\rho_2 - R)^2\right] \tag{S4}$$

where $\rho_1 = \sqrt{y^2 + z^2}$ is the distance from the first fiber (the $x$ axis) and $\rho_2 = \sqrt{(y-\Delta)^2 + z^2}$ is the distance from a second fiber. (If only one fiber is simulated, only the first term is included.) The typical distance of separation between the fiber and the cell center of mass is taken to be $R$, the radius of the cell. This is appropriate if the fiber is, as in our case, much smaller than the cell radius. This energy reflects the fact that if a cell is pulled away from the fiber ($\rho > R$), it must deform – but so must cells compressed close to the fiber ($\rho < R$).

Cell-cell interactions are pairwise,

$$U_{\text{cell-cell}} = \frac{1}{2}\sum_{i \neq j} V(|\mathbf{r}_i - \mathbf{r}_j|) \tag{S5}$$

with an interaction energy

$$V(r) = \begin{pmatrix} \frac{1}{2}\kappa_{\text{cell}}(r-2R)^2 & r < 2R \\ \frac{1}{2}\alpha(r-2R)^2 & r \in [2R, 2R_{\text{adh}}] \\ \frac{1}{2}\alpha(2R_{\text{adh}} - 2R)^2 & r > 2R_{\text{adh}} \end{pmatrix} \tag{S6}$$

Note that overlap begins at separations $r < 2R$, as $R$ is the cell radius, not its diameter. This potential is shown for different values of $\alpha$ in Fig. S5 – larger values of $\alpha$ correspond to a larger adhesion (i.e. a larger force required to rupture one cell from another).

### III. SIMULATION PROTOCOL

We simulate cell-cell collisions on a fiber. To set up each collision, we first simulate the cells without any physical cell-cell interactions or CIL for a time $T_{\text{relax}}$ (setting $\beta = 0, \alpha = 0, \kappa_{\text{cell}} = 0$); this ensures that the cells have polarities $\mathbf{p}_i$ that are sampled according to the steady-state distribution of individual cells – i.e. that this represents a random cell. We then set up the cells to collide, choosing their $x$ separation to be $x_{\text{init}} = 3R$. (We keep their $y$ and $z$ values to be the values generated by the initial evolution without cell-cell interactions). We also choose the cell polarities to be pointing toward one another, so that head-head collisions are the likely outcome. (We do this by ensuring that the cell that has the larger $x$ has a negative $p_x$ and the cell with a smaller $x$ has positive $p_x$. If necessary, we flip the sign of one of the cell's $p_x$ values; this is permitted because in the absence of interactions $p_x \to -p_x$ is a symmetry of our polarity equation.) After this setup, we run the simulation for a time $T_{\text{sim}}$ and track the outcome.

In modeling collisions where one cell has recently divided, we first perform the relaxation as above, but for the cell which has divided, we multiply its polarity along the axis $p_x$ by a factor $M$. We note that the initial polarity $p_x$ for each cell is set by the relaxation process, and so is random; this means that the daughter cell's $p_x$ is not necessarily larger than the other cell's $p_x$, reflecting the large variability in speeds.

We run $N_{\text{it}} = 200$ head-head collisions for each set of parameters and automatically classify the outcomes with the rules:

- **No contact.** If the cells never reach an $r$ separation of $2R_{\text{adh}}$ (i.e. they never interact), we note this as a non-collision; outcome fractions are computed as fractions of collision events.
- **Reversal.** A collision is marked as a reversal if the cells contact, then separate by $x_{\text{init}}$ with the cell at initially lower $x$ remaining at lower $x$
- **Walk-by.** A collision is marked as walk-by if the cells contact, and the $x$ positions of the cells exchange (the cell at smaller $x$ is above the cell at higher $x$). If a trajectory could be counted as either walk-by or reversal (due to multiple collisions), whichever event occurs first is counted.
- **Train/stalling/push.** If the cells have contacted but not separated more than $x_{\text{init}}$, we count the collision as a training event or stalling. (This would also encompass a pushing event.) Because cells in contact will eventually polarize and move as a train, we do not distinguish between these different outcomes in our statistics.

### IV. EFFECT OF CHANGING PARAMETERS ON OUTCOMES

In this section, we take our best fit parameters (Table S3) as our "wild type" and show how the outcomes depend on varying key parameters. We show this for four cases: head-head collisions in the spindle geometry, in the parallel geometry, spindle with a division, and parallel with a division. In all cases, up-regulating the adhesion strongly suppresses walk-past and promotes train formation (Fig. S6a): if cells are not able to separate from one another, they will move together. Another relevant parameter is the degree of attachment between the cell and the fiber(s) – essentially the strength of the spring holding the cell to the surface of the fiber. We find that this primarily plays a role when the cells are attached to two fibers: in the parallel geometry, increasing this strength suppresses walk-by (Fig. S6b), though it has little effect in the single-fiber geometry. This is again intuitively likely: for cells to walk past each other in the parallel geometry, they must move away from their preferred position precisely between the two fibers – and this is suppressed by increasing the adhesion to the fiber. Perhaps surprisingly, the strength of CIL $\beta$ only weakly regulates outcomes, tending to promote training and suppress walk-past (Fig. S6c). This relatively weak dependence on $\beta$ is in part because we have not distinguished in our outcomes between cells that turn around rapidly on contact, and those that take larger times to repolarize.

Walk-past can be strongly increased by having cells with larger polarity noise $\sigma$ – cells with more variable velocities are likelier to walk past each other (Fig. S7a). However, we find no apparent effect from changing the cell's stiffness itself (Fig. S7b), likely reflecting that the cell stiffness found in the fit is relatively high. In addition, we see that our time step does not significantly influence rates (Fig. S7c), showing that the simulations are numerically converged.

### V. PARAMETER FITTING AND VARIATION

#### A. Fixed parameters

We show parameters that are fixed through all our simulations in Tables S1 and S2. We note that these parameters (except the fiber spacing) are also kept constant between spindle and parallel simulations; we have taken a minimal approach and not directly treated changes in, e.g. persistence time or $v_{cg}$ between spindle and parallel. All of the effects of the spindle-parallel geometry change in the simulation is due to the presence of the additional fiber.

#### B. Parameters that are varied

The parameters shown in Table S3 are used in two different ways. First, we generate a plausible range of parameters, and evaluate collision statistics over a sample enclosing that whole range, generating a large number of different plausible parameter sets. We show in the main paper that the main qualitative conclusions are robust to these parameter variations (Figure 5 of the

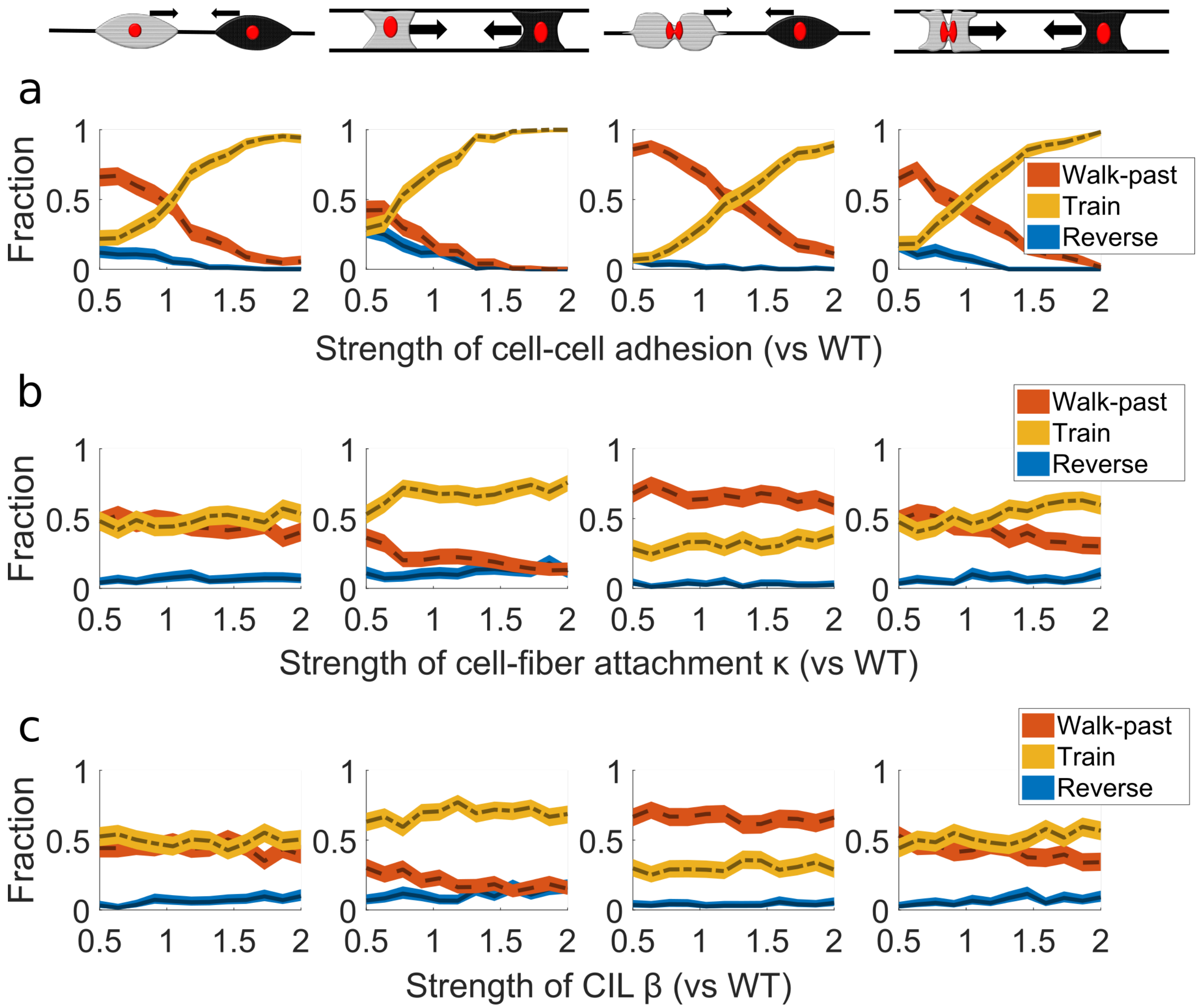


FIG. S6. **Biophysical parameters controlling the relative rates of different outcomes.** The fraction of each outcome is shown as a function of a single parameter being varied relative to its "WT" (best fit to experimental data) value. Shown are a) cell-cell adhesion strength, b) cell-fiber attachment strength, and c) strength of CIL. The four columns show the four collision types – spindle-spindle, parallel-parallel, spindle-daughter, and parallel-daughter. Shaded regions indicate 95% confidence interval, computed via the Goodman method [6] assuming the distribution of outcomes is multinomial. Each point on the lines is computed from 200 individual simulations.

main paper). Secondly, we find a best global fit to the observed experimental outcomes (e.g. spindle-spindle collisions producing 66% walk-past; the comparison is Figure 6 of the main paper). Both the range and the best fit are shown in Table S3.

To generate the range of representative parameters, we used Latin hypercube sampling, so that many combinations of parameters were explored. Because the plausible range of the parameters $\beta$, $\kappa$, and $\kappa_{\text{cell}}$ were large, we chose our sampling to be logarithmically spaced over the plausible range.

To find our best fit, we minimized the mean-square error of the fractions $f_i$ of events between experiment and simulation (using $N_{\text{it}} = 200$ iterations). The minimization was done with Nelder-Mead unconstrained optimization (MATLAB's fminsearch), starting from the minimal error point found in the Latin hypercube sample. We have also put in a small penalty term to changing $\sigma$, which is more closely constrained by the data.

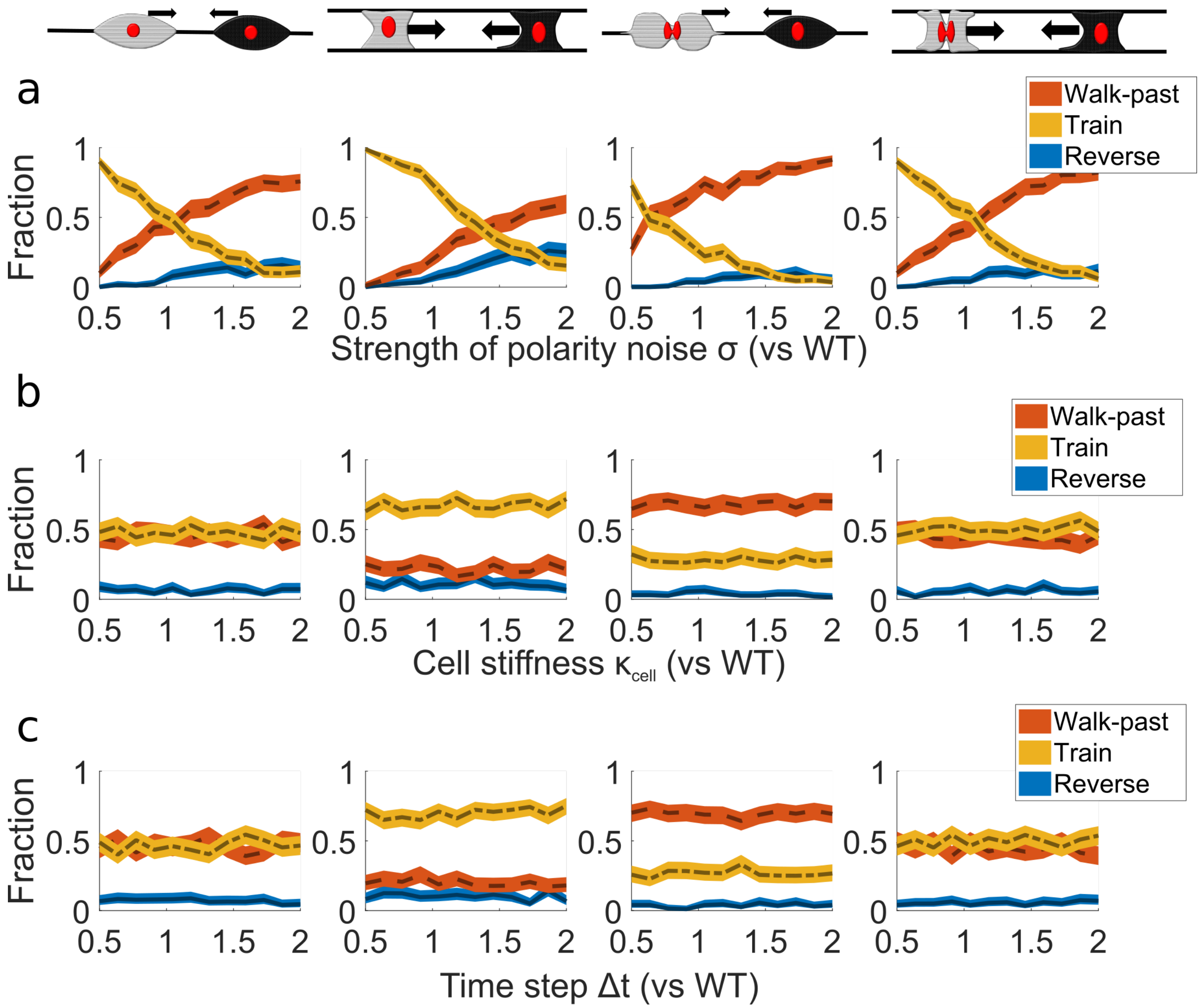


FIG. S7. **More biophysical and computational parameters controlling the relative rates of different outcomes.** The fraction of each outcome is shown as a function of a single parameter being varied relative to its "WT" (best fit to experimental data) value. Shown are a) cell polarity noise $\sigma$, b) cell stiffness $\kappa_{cell}$, and c) the time step $\Delta t$. The four columns show the four collision types – spindle-spindle, parallel-parallel, spindle-daughter, and parallel-daughter. Shaded regions indicate 95% confidence interval, computed via the Goodman method [6]. Each point on the lines is computed from 200 individual simulations.

TABLE S1. Table of fixed biological model parameters

| **Parameter** | **Name** | **Value** | **Units** | **Justification** |
|---|---|---|---|---|
| $\tau$ | Persistence time | 3 | hours | Measured from our experiments showing that cells lose their post-division additional velocity in $\sim 3$ hours. |
| $v_{cg}$ | Contact guidance velocity | 18 | $\mu m$/hr | Estimated from typical time-averaged $x$ cell velocities in our experiment |
| $M$ | Effect of division | 3 | unitless | Rough estimate from experiments; this reflects the increase in a time-averaged speed along the $x$ axis, not merely the instantaneous speed in any direction. |
| $R$ | Cell radius | 25 | $\mu m$ | Rough order of magnitude from experiment. We note that "radius" of a cell here will always be a little misleading because the cells are extremely elongated; we chose this value to be a compromise between the typical cell extensions |
| $R_{\text{adh}}$ | Adhesion radius | 30 | $\mu m$ | Must be larger than cell radius |
| $\Delta$ | Fiber spacing | 50 | $\mu m$ | Chosen to be exactly $2R$ so that one cell fits precisely between fibers |

TABLE S2. Table of parameters related to simulation details

| **Parameter** | **Name** | **Value** | **Units** | **Justification** |
|---|---|---|---|---|
| $\Delta t$ | Time step | 0.01 | hr | Set by convergence |
| $x_{\text{init}}$ | Initial separation | 3 $R$ | $\mu m$ | Set to be relatively small so a high fraction of cells collide. |
| $T_{\text{relax}}$ | Equilibration time | 30 | hr | This is the time for which we simulate each cell in isolation before we set them up to collide. This is set to be much longer than the cell persistence time. |
| $T_{\text{sim}}$ | Simulation time | 15 | hr | Time which is simulated for each collision; if cells have not contacted during this time, no collision is counted. |

TABLE S3. Table of parameters related to simulation details

| Parameter | Name | Plausible range | Best fit | Units | Justification |
|---|---|---|---|---|---|
| $\sigma$ | Noise strength | 8.57-15.91 | 12.21 | $\mu m\,\mathrm{hr}^{-3/2}$ | Range chosen so that typical velocity standard deviation $\sqrt{\sigma^2\tau/2}$ is roughly $0.25$ microns/min, to be consistent with observed variance in velocities; we have included a range of possible variation of 30% around this value, as we suspect it is not well-constrained due to small sample size |
| $\kappa\mu$ | Strength of fiber-cell attachment | 1-50 | 1.81 | $\mathrm{hr}^{-1}$ | $\kappa$ has units of a spring constant, but it only enters into our model in the combination $\kappa\mu$. This range encompasses both values where the cell is tightly attached to the filament and cannot be pulled from it and those where it is more easily stretched from contact with the filament, but rules out parameters where a single cell will spontaneously move away from contact with the wires. *Note: To interpret this and other adhesion strengths/stiffnesses with these units, it's useful to think that $\kappa\mu = 10/hr$ means that under a deformation of $1$ micron, the responding velocity of the cell is $\kappa\mu \times 1 micron = 10 micron/hr$.* |
| $\kappa_{\mathrm{cell}}\mu$ | Cell stiffness | 10-50 | 23.7 | $\mathrm{hr}^{-1}$ | $\kappa_{\mathrm{cell}}$ is a spring constant, but it only enters into our model in the combination $\kappa_{\mathrm{cell}}\mu$. This range of values is set so that the cell-cell repulsion is strong enough that cell overlaps are not common – this requires $\kappa_{\mathrm{cell}}\mu \sim O(10)\mathrm{hr}^{-1}$ |
| $\alpha\mu$ | Strength of cell-cell adhesion | 2.2-4 | 3.11 | $\mathrm{hr}^{-1}$ | $\alpha$ is a spring constant, entering into the model only as $\alpha\mu$. This range is chosen because if $\alpha\mu$ is much outside this range, either cells will never be able to adhere or they will never be able to separate. |
| $\beta$ | CIL strength | 8-40 | 15.4 | $\mathrm{hr}^{-1}$ | We know that, in order to create a relevant effect, we must have $\beta\tau$ – the maximum velocity that can be induced by CIL – on the order of cell velocities ($20\,\mathrm{microns/hr}$). We chose the low end of this range to ensure that cells in contact did repolarize – at the high end, the induced velocities become unreasonable. |
